# Comparative Modeling of Transcranial Magnetic and Electric Stimulation in Mouse, Monkey, and Human

**DOI:** 10.1101/442426

**Authors:** Ivan Alekseichuk, Kathleen Mantell, Sina Shirinpour, Alexander Opitz

## Abstract

Transcranial magnetic stimulation (TMS) and transcranial electric stimulation (TES) are increasingly popular methods to noninvasively affect brain activity. However, their mechanism of action and dose-response characteristics remain under active investigation. Translational studies in animals play a pivotal role in these efforts due to a larger neuroscientific toolset enabled by invasive recordings. In order to translate knowledge gained in animal studies to humans, it is crucial to generate comparable stimulation conditions with respect to the induced electric field in the brain. Here, we conduct a finite element method (FEM) modeling study of TMS and TES electric fields in a mouse, capuchin monkey, and human model. We systematically evaluate the induced electric fields and analyze their relationship to head and brain anatomy. We find that with increasing head size, TMS-induced electric field strength first increases and then decreases according to a two-term exponential function. TES-induced electric field strength strongly decreases from smaller to larger specimen with up to 100x fold differences across species. Our results can serve as a basis to compare and match stimulation parameters across studies in animals and humans.

**HIGHLIGHTS:**

- Translational research in brain stimulation should account for large differences in induced electric fields in different organisms
- We simulate TMS and TES electric fields using anatomically realistic finite element models in three species: mouse, monkey, and human
- TMS with a 70 mm figure-8 coil creates an approximately 2-times weaker electric field in a mouse brain than in monkey and human brains, where electric field strength is comparable
- Two-electrode TES creates an approximately 100-times stronger electric field in a mouse brain and 3.5-times stronger electric field in a monkey brain than in a human brain

## 1. Introduction

Non-invasive brain stimulation (NIBS) is a promising method to study causality of brain-behavior relationships in humans as well as for clinical research in neurological and psychiatric disorders (Polanía et al., 2018). Two main methods are currently used: transcranial magnetic stimulation (TMS) and transcranial electric stimulation (TES). TMS affects neural tissue by inducing a short-lasting electric field at sub-or suprathreshold intensities via electromagnetic induction (Valero-Cabré et al., 2017). TES generates a long-lasting subthreshold electric field that aims to modify spike timing by directly applying electric currents to the scalp (Paulus et al., 2016). The induced electric field in the brain is the main actor for both TMS and TES effects. However, the electric field is also the most difficult feature to predict as it depends not only on controllable factors, such as current intensity and coil or electrode locations, but also on the individual head anatomy and tissue biophysics (Alekseichuk et al., 2018; Datta et al., 2009; Laakso et al., 2015; Miranda et al., 2013; Opitz et al., 2017, 2015, 2011; Peterchev et al., 2012; Thielscher et al., 2011; Wagner et al., 2014).

Stimulation-induced electric fields are difficult to directly assess in humans except in cases of intracranial measurements in surgical epilepsy patients. Thus, modeling approaches are most often used to study NIBS electric field distributions (Miranda et al., 2018). While computational models are clearly useful to guide stimulation protocols and to ensure target engagement (Huang et al., 2017; Opitz et al., 2018, 2016), they still cannot predict the physiological outcome of NIBS studies. This is due to the missing link between the biophysics of stimulation, i.e. electric fields, and the resulting physiological effects. Animal models are crucial to close this knowledge gap because they allow simultaneous measurement of both the biophysics and physiology of NIBS using invasive recordings.

Invasive studies in animal models offer a larger neuroscientific toolset with a higher spatial precision than noninvasive evaluations in humans. Thus, animal work is increasingly used to dissect NIBS mechanisms (Kar et al., 2017; Krause et al., 2017; Vöröslakos et al., 2018). However, the translation of results from the animal literature to humans is challenging because it is unclear how to transfer stimulation parameters and dose regimes to achieve comparable conditions. To date, TES research in small rodents predominantly utilizes currents at 100 to 200 μA peak-to-baseline (Grossman et al., 2017; Liebetanz et al., 2006; Monai et al., 2016; Pedron et al., 2014), while some work uses weaker inputs of 20-100 μA (Faraji et al., 2013; Wachter et al., 2011) or higher than 200 μA (Cambiaghi et al., 2011; Takano et al., 2011). TES in non-human primates typically operates at the intensity of 1-2 mA (Kar et al., 2017; Krause et al., 2017). The same intensity is most common in human studies and clinical applications (Antal et al., 2017; Paulus et al., 2016). For TMS, the stimulation intensities used are in the same range for animal (Hoppenrath and Funke, 2013; Mueller et al., 2014; Pasley et al., 2009) and human studies (Rossi et al., 2009). Further, many animal studies use smaller TMS coils, which result in more focal induced electric fields to compensate for the smaller head size (Deng et al., 2013). There is an implicit assumption that the NIBS dose regimens across species are comparable, yet there is limited evidence and a lack of systematic evaluations.

Here, for the first time, we conduct a systematic comparison of electric fields in the brain during (i) TMS with either a 70 or 25 mm figure-8 coil and (ii) two-electrode TES using realistic FEM models of a mouse, monkey, and human. For both methods, we consider multiple coil/electrode positions and identify relationships between the electric field properties and the properties of the head volume conductor.

## 2. Materials and methods

### 2.1. General modeling framework

A realistic whole-body mouse model and head models of a capuchin monkey and a human were created from structural MRI images as described below following the SimNIBS framework (Nielsen et al., 2018; Windhoff et al., 2013). In addition, we generated a range of six-layer spherical models with varying radii to study the effect of head size on NIBS electric fields in a simplified scenario. For all models we simulated electric fields for transcranial magnetic stimulation (TMS) and transcranial electric stimulation (TES). If not stated otherwise, options were set to the SimNIBS defaults with the following isotropic conductivities: skin and soft tissue (σ = 0.465 S/m), bone (σ = 0.01 S/m), eyes (σ;= 0.5 S/m), CSF (σ = 1.654 S/m), grey matter (σ = 0.275 S/m), and white matter (σ = 0.126 S/m; not included in the mouse model). Resulting electric fields in the brain (grey matter) were analyzed and compared in MATLAB.

### 2.2. FEM models

#### Mouse

We created a FEM model from a segmented individual anatomical atlas of a normal adult male nude mouse “Digimouse” (Dogdas et al., 2007). The atlas was derived from X-ray CT and cryosection images. X-ray CT images were acquired in two bed positions using an Imtek microCAT system (Imtek, Knoxville, TN) and reconstructed with 0.1×0.1×0.1 mm resolution. Cryosections were cut at a thickness of 50 μm with an in-plane resolution of 38.8×38.8 μm. The data were co-registered and resampled on an isotropic 0.1 mm grid. Then, images were segmented for soft tissues, skeleton, and multiple internal organs. Here, we considered the following structures: soft tissues, bones, eyes, and the whole brain including the cerebellum, medulla, and olfactory bulbs. We manually refined the brain surface to correct remaining small defects and introduced a layer of CSF using FSL (Jenkinson et al., 2012). 3D surfaces were created from the refined atlas using FreeSurfer (Fischl, 2012), and further optimized with MeshFix (Attene, 2010). A tetrahedral based FEM model was generated employing adaptive meshing in Gmsh (Geuzaine and Remacle, 2009) with GM resolved at a higher numerical resolution. The whole-body model comprises ~ 5.8 million tetrahedral elements.

**Figure 1.**
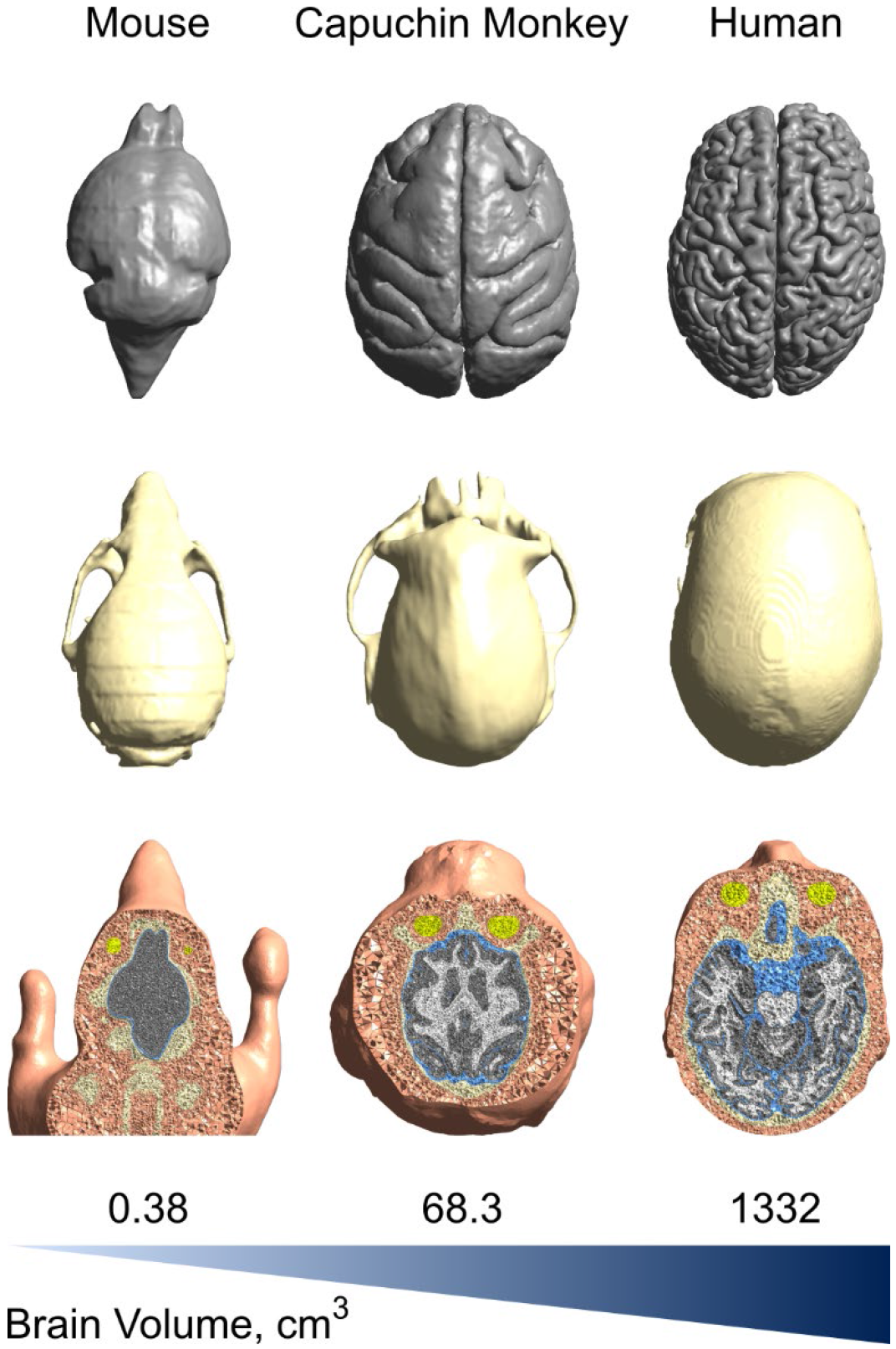
Anatomically accurate FEM models of mouse, monkey, and human. The following tissues are considered: skin and soft tissues, skull and bones, eyes, CSF, grey matter, and white matter. On the top row are the brain surfaces, middle row – skull surfaces, bottom row – horizontal cut of FEMs.

#### Monkey

We used an individual FEM model of a normal adult male capuchin monkey (from Alekseichuk et al., 2018). In short, structural MR imaging (T1 and T2) was performed at the Nathan Kline Institute for Psychiatric Research, USA with approval of the local Institutional Animal Care and Use Committee. Anatomical images were segmented for the soft tissues, eyes, skull, CSF, white and grey matter using Freesurfer and ITKSnap (Yushkevich et al., 2006). Tissue surfaces were created with FreeSurfer and optimized using MeshFix. The FEM head model (~ 4.2 million tetrahedral elements) was generated using Gmsh.

#### Human

We utilized the individual head model of the normal adult male human “Ernie” that is included in SimNIBS 2.1 (Nielsen et al., 2018). T1-weighted and T2-weighted MR images were collected using a 3T scanner (Phillips Achieva™) with a 32-channel head coil at the Copenhagen University Hospital Hvidovre, Denmark. The Ethics Committee of the Capital Region of Denmark approved the MR scans. A T1-weighted contrast was acquired with the following parameters: 3D TFE, TR/TI/TE = 6.9/1000/3.3 ms; TFE factor = 243, 2600ms; flip angle = 8°; 208 sagittal slices; matrix = 256×256; voxel size = 1×1×1 mm^3^. T2-weighted image: 3D TSE, TR/TE = 2500/250 ms; flip angle = 90°; 208 sagittal slices; matrix = 244×244; voxel size = 1×1×1 mm^3^. The facial region of the images was depersonalized. The following tissues were considered: skin and soft tissues, eyes, skull, CSF, grey matter, and white matter. The model was generated with the SimNIBS routine ‘headreco’, which also utilizes Gmsh for adaptive meshing. The final head model comprises ~ 4.2 million tetrahedral elements.

**Table 1.** Summary of the head models. The brain size is given in the following projections: rostral-caudal (RC), left-right (LR), and dorsal-ventral (DV).

| Model | Brain Size, cm |  |  | Brain Volume, cm<sup>3</sup> | Head Volume, cm<sup>3</sup> |
| --- | --- | --- | --- | --- | --- |
|  | RC | LR | DV |  |  |
| Mouse | 1.71 | 0.94 | 0.72 | 0.38 | 3.10 |
| Monkey | 6.60 | 5.50 | 5.00 | 68.31 | 472.66 |
| Human | 18.55 | 13.60 | 12.95 | 1331.87 | 5270.54 |

#### Spherical models

We created 13 six-layer spherical models with an outer radius of 0.5 cm, and 1 to 12 cm with 1 cm steps. Each model was linearly scaled from the standard spherical model as included in SimNIBS 2.1 and originally created in Gmsh with the following layers: “ventricles with CSF” (r = 25 mm, σ = 1.654 S/m), “white matter” (r = 75 mm, σ = 0.126 S/m), “grey matter” (r = 80 mm, σ = 0.275 S/m), “CSF” (r = 83 mm, σ = 1.654 S/m), “skull” (r = 89 mm, σ = 0.01 S/m), and “skin” (r = 95 mm, σ = 0.465 S/m). Each model has ~ 480 thousand tetrahedral elements. To ensure that found results for differing radii were not due to differences in tetrahedral element size, we also created high resolution spherical models (~ 8 million elements) for r = 8 to 12 cm.

### 2.3. Transcranial magnetic stimulation (TMS)

We simulated the electric field for 70 mm and 25 mm figure-8 coils (Thielscher and Kammer, 2004). The coils were positioned over the left central brain region, which corresponds to the primary motor cortex in humans. We simulated three different orientations: 45° to the rostral-caudal axis (towards caudal medial end, anterior-posterior with respect to the motor cortex), 90° to rostral-caudal axis (towards medial end), and −45° to rostral-caudal axis (towards caudal lateral end). The coil center was placed 4 mm above the skin surface. The input intensity *dI/dt* was 100 A/μs, in a range commonly used in human experiments (Rossi et al., 2009). Overall, 18 simulations (3 FEM models × 3 coil orientations × 2 coil sizes) were performed using SimNIBS 2.1, which uses the GetDP solver (Geuzaine, 2007).

Using the same parameters, we also investigated the induced electric fields in the 13 six-layer spherical models. Only one coil position was used per sphere due to the spherical symmetry. We further performed a control simulation using high-resolution spheres to ensure that found results are not due to differences in the size of tetrahedral elements.

### 2.4. Transcranial electric stimulation (TES)

We estimated the electric field for three different two-electrode montages. The montages aimed to maximize the distance between the electrodes on the head within a given anatomical axis to maximize the electric field strength in the brain. The following montages were modeled: 1) Rostral-caudal montage: one electrode over the medial rostral area (“forehead”) and another over the medial caudal area (“occiput”). 2) Left-right montage: electrodes over the left and right temporal areas. 3) Dorsal-ventral montage: one electrode over the left central area (“motor cortex”) and another over the right shoulder/neck areas. For the mouse model only, an additional dorsal-abdominal montage was simulated; the electrodes were located over the left central brain area and the central abdominal region. Stimulation electrodes were modeled as 2 mm thick rubber (σ = 29.4 S/m) circles with the diameter scaled according to the head size: 3 mm for the mouse model, 15 mm for the monkey model, and 36 mm for the human model. The current to surface area at the electrode-skin interface was 140.85 A/m^2^ for the 3 mm electrode, 5.65 A/m^2^ for the 15 mm electrode, and 0.98 A/m^2^ for the 36 mm electrode. In addition, for the human model we performed simulations for 3 mm and 15 mm stimulation electrodes. The stimulation intensity *I* was set to 1 mA, in line with the human experimental literature (Antal et al., 2017; Grossman et al., 2018). Overall, 16 simulations (3 FEM models × 3 electrode montages + 1 extra montage for the mouse model + 2 extra electrode sizes for the human model × 3 electrode montages) were performed using SimNIBS 2.1.

Using the same computational pipeline, we also estimated the electric fields for the 13 six-layer spherical models. Two round stimulation electrodes (r_electrode_ = 0.1 × r_sphere_) were placed on opposite ends of the sphere. In addition, for the “human head size” sphere (r_sphere_ = 10 cm), all electrode sizes with d = 2 to 30 mm were simulated.

### 2.5. FEM analysis

To quantify the results of TMS and TES simulations, we estimated three main parameters per simulation: robust maximum of the electric field strength (E_max_), which corresponds to the 99.9th percentile of the electric field strength; median of the electric field strength (E_median_); and affected area (L_½max_), which corresponds to the volume or surface where the electric field strength is equal or greater than the half-maximum for the given simulation. We evaluated these parameters in the brain (grey matter) volume and on the brain (grey matter) surface. In addition, tangential and perpendicular/radial components of the induced electric field were separated at the brain surface level and evaluated individually.

We used two-sample t-tests to compare the simulation results in mouse and monkey versus the ones found in the human. We further evaluated the relationship between the parameters E_max_, E_median_, L_½max_ and the total head volume in both the anatomically realistic and spherical FEM models using the MATLAB Curve Fitting Toolbox. We considered a set of plausible functions employing the Levenberg-Marquardt algorithm with a robust nonlinear least squares fitting method. Goodness-of-fit metric, namely adjusted R-squared (R^2^_adj_), is reported for the identified best fit.

## 3. Results

### 3.1. Transcranial magnetic stimulation (TMS)

TMS-induced electric fields were simulated for three coil positions and two coil sizes (70 mm and 25 mm figure-8 coils) in the mouse, capuchin monkey, and human FEM models. All simulations were performed for the same input intensity *dI/dt* of 100 A/μs. Resulting electric field strengths in grey matter are highly comparable between coil positions (Fig. 2A, B), but vary significantly for different coil sizes and head models (Tables A1, A2).

For the 70 mm figure-8 coil, the average robust maximum electric field (E_max_) across the grey matter volume is 56.6 mV/mm for the mouse, 126.4 mV/mm for the monkey, and 126.8 mV/mm for the human (cross-species ratio is 0.45:1:1). A two-sample t-test indicates significant differences between the mouse and human electric field strengths (p = 4.8×10^−5^, t_4_ = 18.72), but not between the monkey and human (p = 0.92, t_4_ = 0.11). The distribution of the electric field on the brain surface across species is shown in Fig. 2C (for other coil positions see Fig. S1). The relative volume of stimulation (L_½max_) also varies significantly: on average 35.8% of brain volume in the mouse, 8.8% in the monkey, and 2.3% in the human is affected (ratio 15.57:3.83:1). Both mouse L_½max_ values (p = 2.1 ×10^−4^, t_4_ = 12.8) and monkey L_½max_ values (p = 1.3×10^−4^, t_4_ = 14.47) are statistically different from the human L_½max_ values.

For the 25 mm figure-8 coil, the grey matter volume average E_max_ is 81.3 mV/mm, 113.4 mV/mm, and 86.5 mV/mm for mouse, monkey, and human, respectively (ratio 0.94:1.31:1). These values are comparable between the mouse and human models (p = 0.25, t_4_ = 1.36), but significantly higher in the monkey model (p = 0.002, t_4_ = 7.17). Average L_½max_ is 33.5%, 6%, and 1.3% for mouse, monkey, and human, respectively (ratio 25.77:4.62:1). The differences are significant both for the mouse model (p = 1.46×10^−4^, t_4_ = 14.11) and monkey model (p = 1.45×10^−4^, t_4_ = 14.14) relative to the human model.

**Figure 2.**
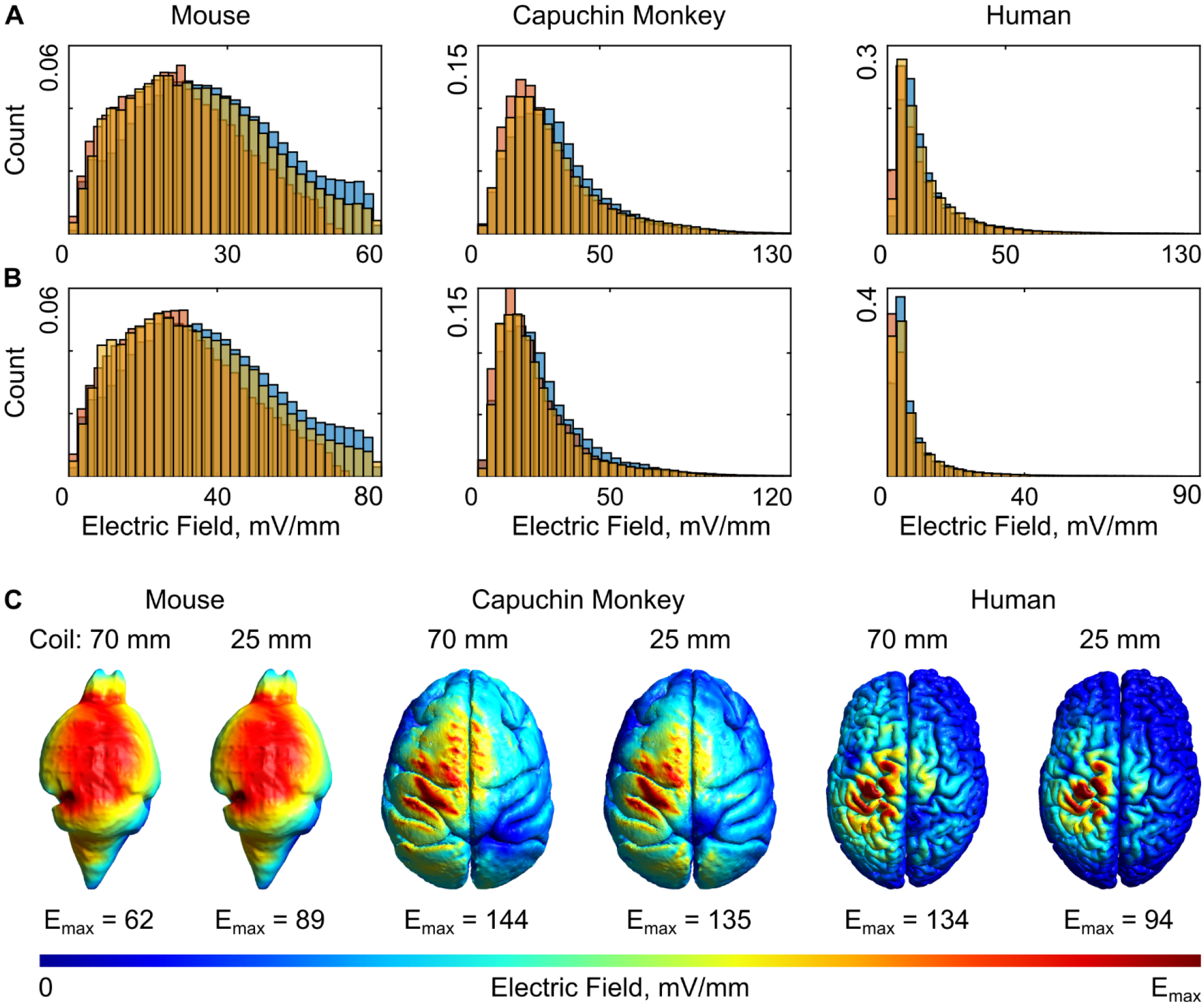
Distribution of the electric field strength in the grey matter volume for a 70 mm (A) and 25 mm (B) figure-8 coil. Blue color corresponds to the caudal-medial or CM coil orientation, red color-medial or M, and yellow–caudal-lateral or CL coil orientation. (C) Normalized electric fields on the brain surfaces for the CM orientation. See other montages in Figures S1, S2.

Comparing the small 25 mm to the standard 70 mm double coil, E_max_ is higher in the mouse (p = 0.003, t_4_ = 6.4), yet lower in the monkey (p = 0.01, t_4_ = 4.34) and the human (p = 4.1×10^−4^, t_4_ = 10.87). L_½max_ is comparable between the coils in the mouse (p = 0.54, t_4_ = 0.67), but in the monkey and human L_½max_ is lower for the 25 mm coil (monkey: p = 0.006, t_4_ = 5.33; human: p = 0.01, t_4_ = 4.25).

**Figure 3.**
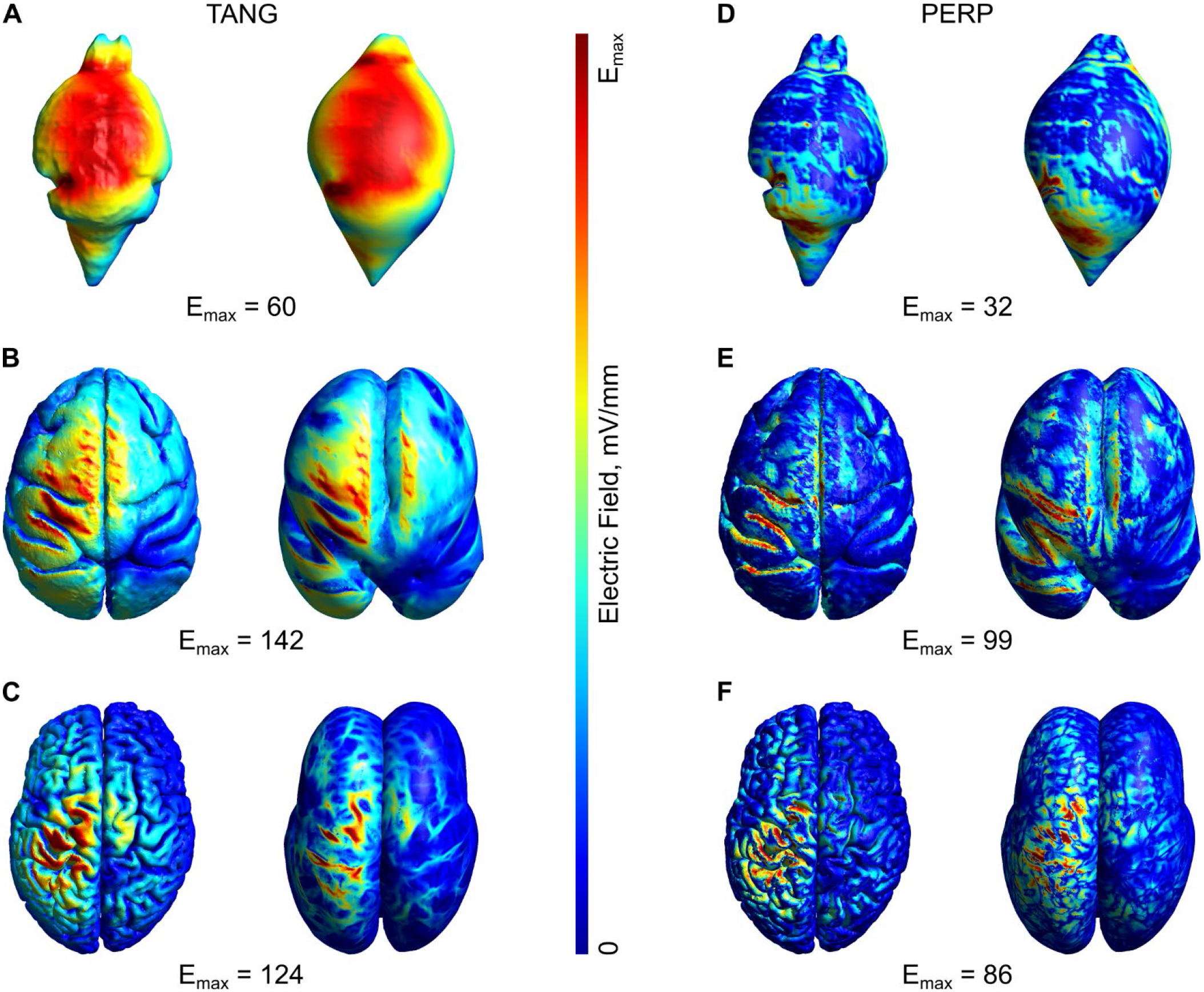
Distribution of the tangential (A-C) and perpendicular (D-F) components of the electric field for the 70 mm figure-8 coil oriented caudal-medial for mouse (top row), monkey (middle row), and human (bottom row). For each figure in the panel, both the anatomical realistic surface (left) and the inflated surface (right) is shown.

Considering that the physiological effects of the electric field depend on its orientation in the brain (Balslev et al., 2007; Brasil-Neto et al., 1992; Opitz et al., 2013; Richter et al., 2013), we separated the electric field into tangential and perpendicular components (Fig. 3 and Table A1). We then further analyzed the spatial distributions of tangential and perpendicular components.

On average, for the 70 mm figure-8 coil, the ratio of the median magnitude (E_median_) of the tangential and perpendicular field components is 2.86, 1.79, and 1.72 in mouse, monkey, and human, respectively. These ratios relate to each other as 1.66:1.04:1. The same pattern exists for the 25 mm figure-8 coil: the ratio of E_median_ of the tangential to perpendicular components is 2.75, 1.77, and 1.71 in mouse, monkey, and human, respectively (relate as 1.61:1.04:1).

A summary of all data is depicted in Figure 4A, B, C. As shown above, TMS-induced electric fields in the brain are getting stronger from the mouse to monkey and human for the 70 mm figure-8 coil. For the 25 mm figure-8 coil, first an increase from mouse to monkey is visible and then a decrease from monkey to human. Regarding relative affected volume L_½max_, it decreases from mouse to monkey to human for both coil sizes.

To analyze how brain/head size affects the electric fields in a simplified model, we computed the TMS-induced electric field in ideal six-layer spherical models of different sizes (r = 0.5 to 12 cm). This size range includes approximations of animal models, such as small rodents (r = 0.5-1 cm) and nonhuman primates (r = 4-6 cm), and humans (r = 9-11 cm). Here, we found a two-term exponential relationship between the electric field strength and head dimensions: E_max_ first increases and later decreases with increasing head size (R^2^_adj_ ≈ 1; Fig. S4A). We confirmed that the found results are not due to differences in the size of tetrahedral elements by re-running the simulations for the large spheres at higher resolution.

Further, given the complex relationship of the electric field strength and brain volume in our numerical simulations, we independently implemented an analytical approach. For this, we calculated the induced electric field in a spherical model for a simple figure-8 wire loop using an analytical formulation (originally derived by Eaton, 1992; further details in Appendix B). Our analytical approach also arrives at a two-term exponential relationship between the electric field and brain size (Fig. S4B) with first increasing and then decreasing field strength. Importantly, this relationship is preserved even after disregarding the factor of coil-to-brain distance, which otherwise increases with the overall head dimensions. To examine this, we simulated a single-layer spherical model where the distance between the TMS coil and the volume of interest is kept identical for every sphere size. Both modeling (Fig. S4C) and analytical solution (Fig. S4D) still show a two-term exponential relationship.

**Figure 4.**
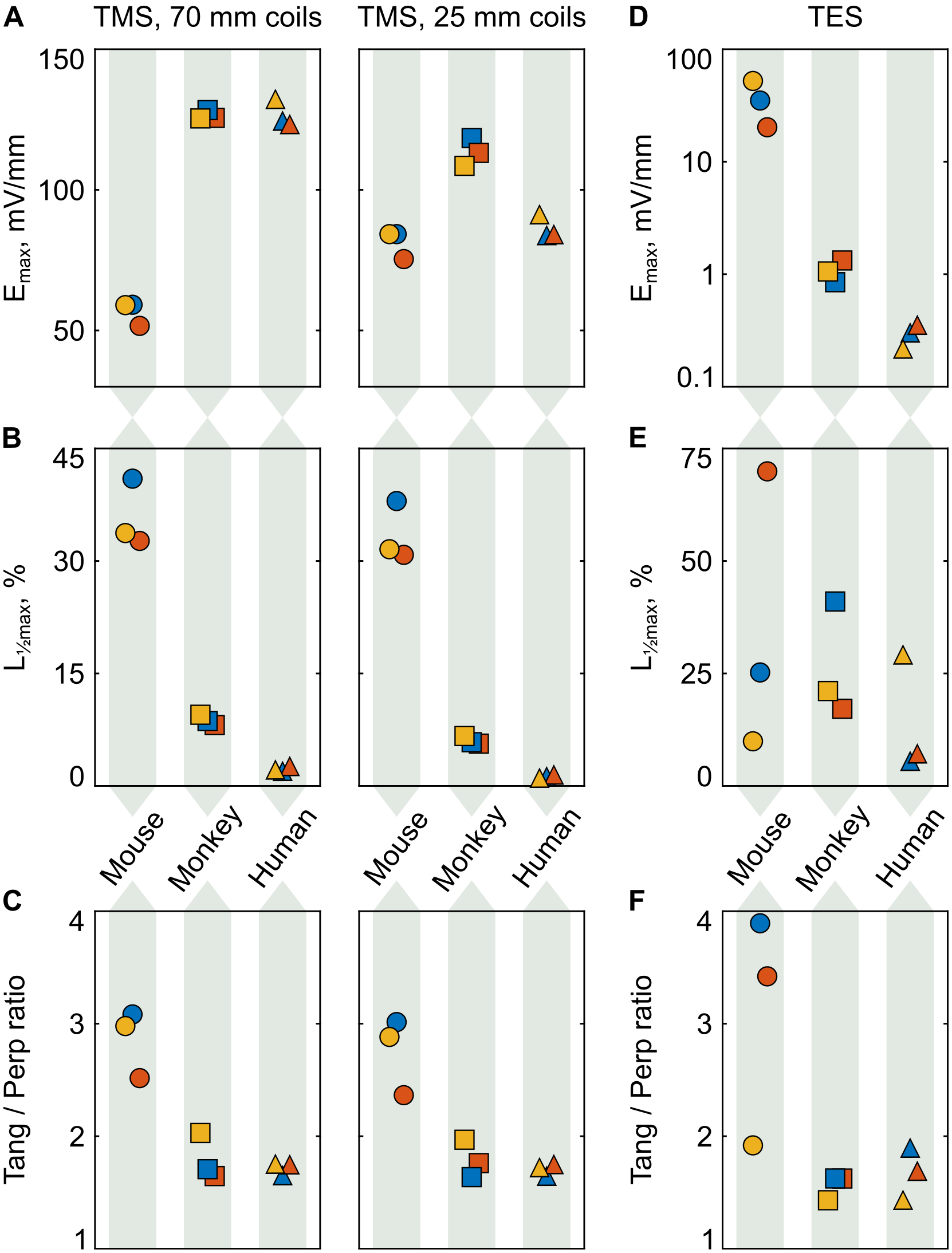
(A-C) TMS summary statistics across species. Color encodes the coil orientation: blue – caudal– medial, red – medial, and yellow – caudal-lateral. (D-F) TES summary statistics across species. Color encodes the electrode montage: blue – rostral-caudal, red – left-right, and yellow – dorsal-ventral. The shape of the data points indicates the species: round – mouse, square – monkey, triangle – human. The top row depicts the robust maximum of the electric field (E_max_), the middle row – the affected brain volume (L_½max_), and the bottom row-the ratio of the medians of the tangential to perpendicular field components.

With increasing head volume, the TMS-induced electric field in the brain first gets stronger and then weaker following an exponential function. The deflection point occurs earlier for a smaller, 25 mm figure-8 coil than for the bigger 70 mm figure-8 coil. For the latter, both monkey and human head sizes are near the peak of the function.

### 3.2. Transcranial electric stimulation (TES)

We modeled TES for three two-electrode montages in a mouse, capuchin monkey, and human model. The stimulation intensity was set to the same level of 1 mA for every simulation. We further modelled an additional electrode montage specific for rodent studies with one electrode located over the head and one over the abdominal area. The results for this montage are highly similar to the dorsal-ventral montage, so it was not included in the group statistics below. All results are shown in Tables A3 and A4.

**Figure 5.**
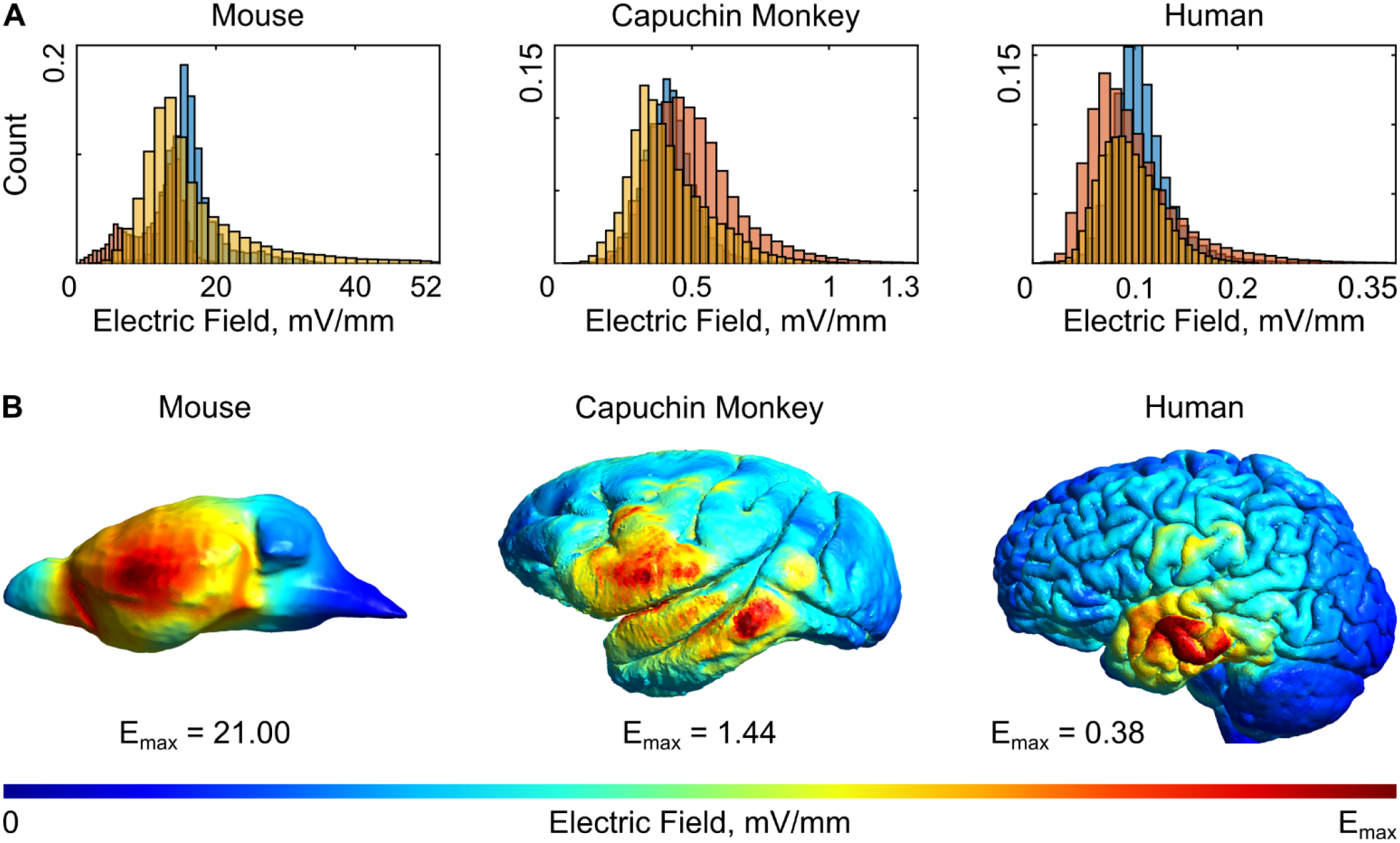
(A) Distribution of the electric field magnitude in the grey matter volume during TES with the following electrode montages: blue color – rostral-caudal, red – left-right (L-R), and yellow – dorsal-ventral. (B) Normalized electric fields on the brain surface for the L-R montage. See the results for the other montages in Figure S3.

First, we compared electric fields across electrode montages. Here we found that the robust electric field strength maximum (E_max_) in the brain shows a wide disparity between montages: 34.98, 20.24, and 52.18 mV/mm for the mouse (mean = 35.8 mV/mm); 0.85, 1.32, and 1.06 for the monkey (mean = 1.08 mV/mm); and 0.3, 0.35, and 0.22 for the human model (mean = 0.29 mV/mm). Average ratio across species is 123.45:3.72:1 (see Fig. 5 and Fig S3). Two-sided t-tests indicate significant differences both for the mouse (p = 0.02, t_4_ = 3.85) and monkey electric field maxima (p = 5.1 × 10^−3^, t_4_ = 5.58) relative to the human.

Considering the relative brain volume of stimulation (L_½max_), we found 25.23%, 69.91%, and 9.93% for the mouse (mean = 35.02%); 41%, 17.15%, 21.1% for the monkey (mean = 26.42%), and 5.58%, 7.23%, and 29.21% for the human (mean = 14%). The ratio of these values is 2.5:1.89:1. However, high variability between the different electrode montages resulted in no statistically significant differences across species (t-test p > 0.05).

**Figure 6.**
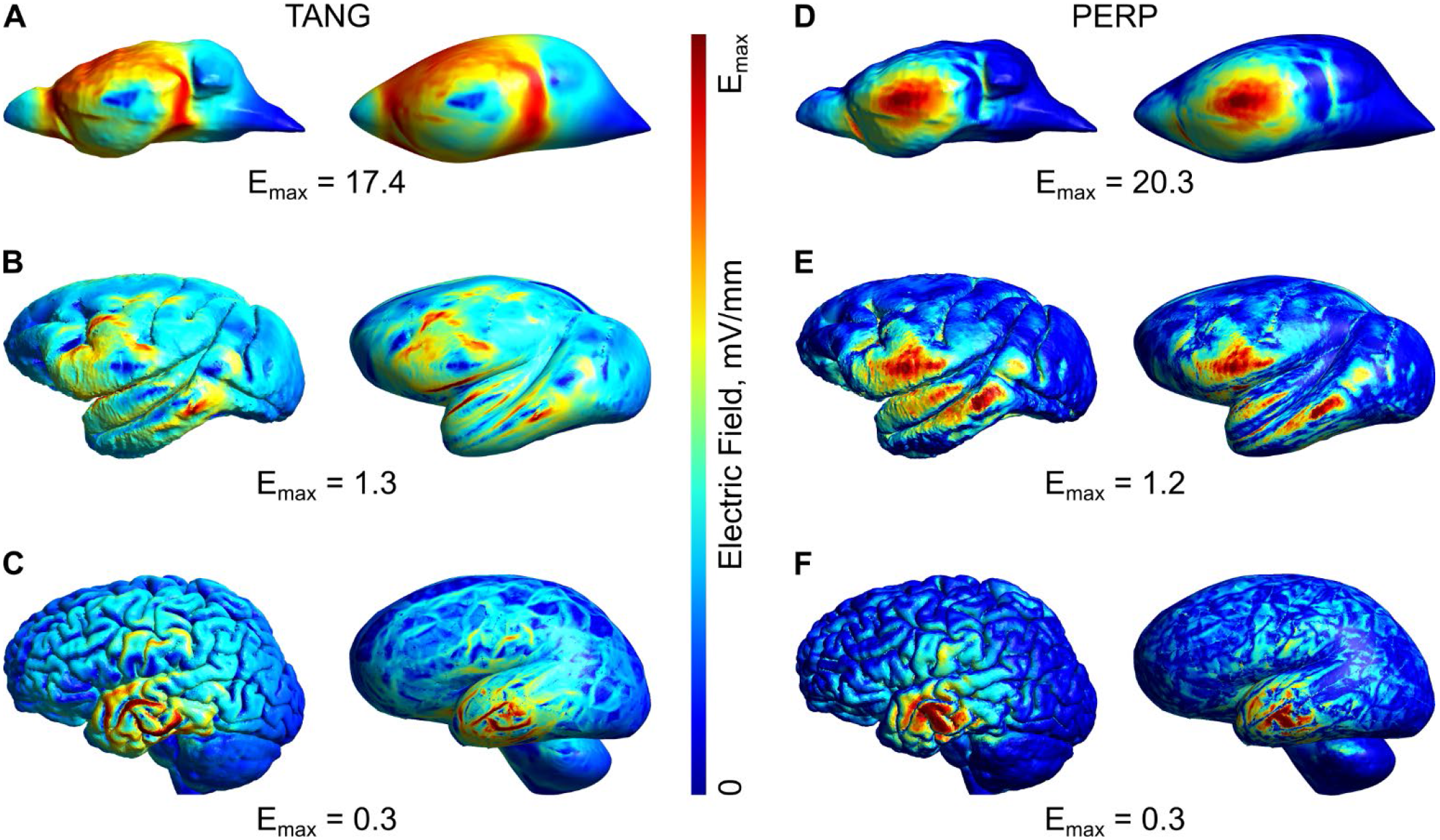
Distribution of the tangential (A-C) and perpendicular (D-F) components of the electric field on the brain surface due to TES with the left-right electrode montage. On the top row is mouse, middle row – monkey, and bottom row – human. For each figure in the panel, normal surface is on the left and its inflated version is on the right.

In addition, we evaluated the tangential and perpendicular components of the electric field in a separate analysis (Fig. 6). The ratios of the median magnitude (E_median_) of the tangential to perpendicular electric field components are 3.89, 3.42, and 1.92 in the mouse (mean = 3.08); 1.63, 1.64, and 1.42 in the monkey (mean = 1.56); 2, 1.5, and 1.5 in the human (mean = 1.67). Mean values relate as 1.84:0.93:1.

The summary of all TES data is shown in Figure 4D, E, F. Unlike TMS, E_max_ estimates for TES are decreasing from smaller to larger organisms. At the same time, the relative affected volume L_½max_ varies greatly due to the large variability across electrode montages.

We also simulated TES electric fields in ideal spherical head models. We simulated 13 six-layer spheres with radii r = 0.5 cm and from 1 to 12 cm with two electrodes attached on opposite sites (Fig. S5A). We found that the maximum electric field strength E_max_ exponentially decreased with increasing radius (R^2^_adj_ ≈ 1).

Considering the differences in TES applications across animal and human studies, one apparent distinction is the electrode sizes. Naturally, electrodes to be used for mouse stimulation are much smaller than for those in humans. Here, we simulated round electrodes with d = 3 mm for the mouse, 15 mm for the monkey, and 36 mm in humans. Given the same stimulation intensity *I* of 1 mA, the ratio of current to surface area *I/A* at the electrode-skin interface was 140.85, 5.65, and 0.98 A/m^2^, respectively. To investigate the role of different electrode sizes, we conducted an additional series of simulations in the human head model using all three above-mentioned electrode sizes (Table A4). For the left-right electrode montage, the maximum electric field E_max_ in the grey matter volume had only a weak dependency of the *I/A* ratio: E_max_ = 0.43, 0.40, and 0.35 mV/mm for *I/A* = 140.85, 5.65, and 0.98 A/m^2^, respectively (Fig. 7B, linear R^2^_adj_ = 0.29), with slightly higher fields for a smaller surface area. The two other montages, rostral-caudal and dorsal-ventral, did not show such a trend (Fig. 7A, C). We further evaluated the effect of *I/A* ratio in the spherical model with r = 10 cm and stimulation electrodes size ranging from d = 2 to 30 mm (Fig. S5B, C). We found a significant linear decrease in E_max_ with increasing electrode size (R^2^_adj_ = 0.89, slope β_1_ = −0.004). However, the slope of the regression is small with values ranging from 0.234 mV/mm for 2 mm electrodes to 0.184 mV/mm (−21.37%) for 30 mm electrodes. Altogether, the difference in electrode sizes for mouse, monkey, and human had only a small effect on the found results.

**Figure 7.**
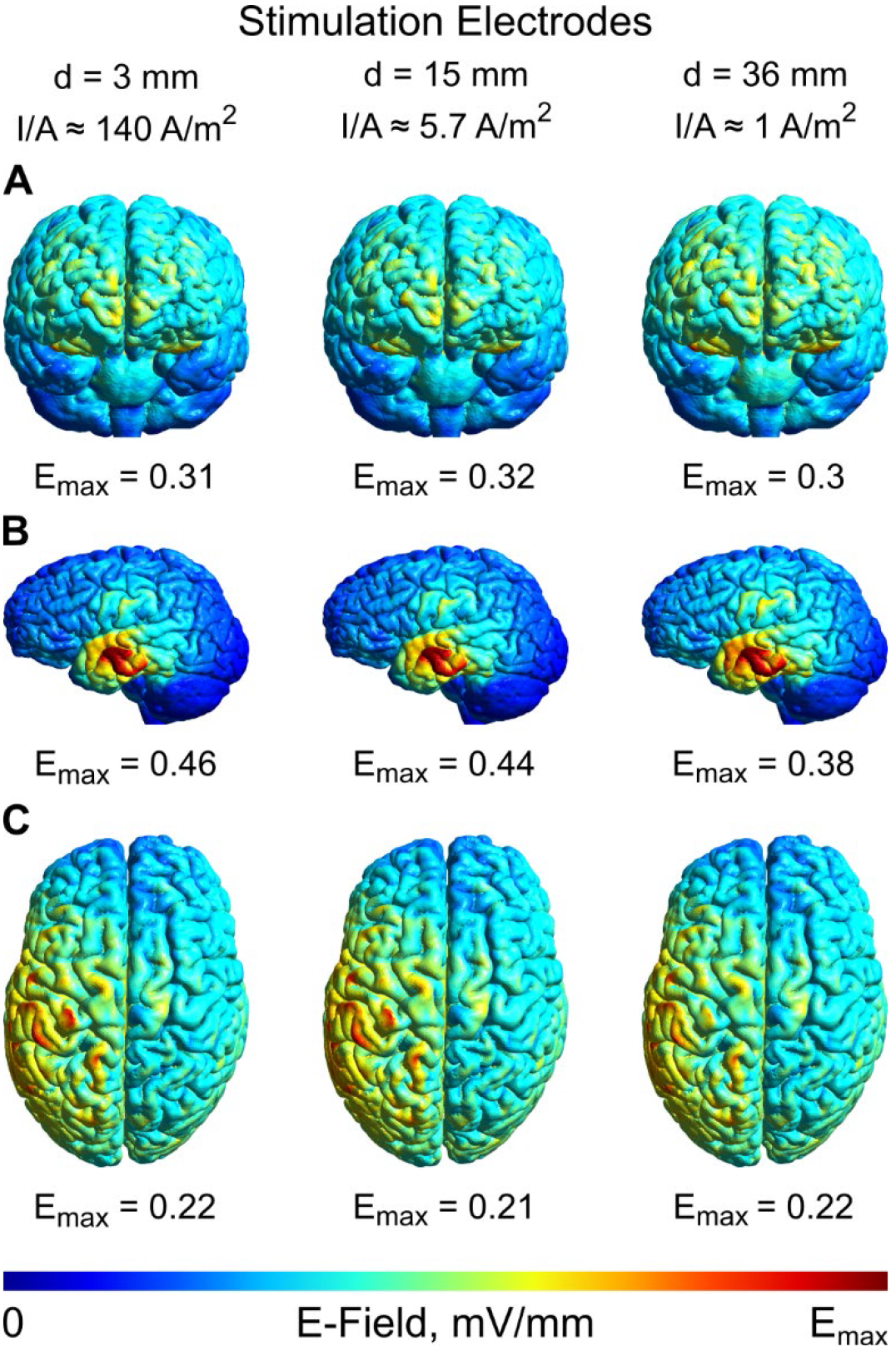
TES electric fields in gray matter for different sizes of stimulation electrodes (d = 36 mm, 15 mm, and 3 mm). (A) Frontal view for the rostral-caudal electrode montage. (B) Lateral view for the left-right montage. (C) Transverse view for the dorsal-ventral montage.

## 4. Discussion

Here, we systematically evaluated the TMS- and TES-induced electric fields in realistic mouse, monkey, and human FEM models. We directly compared electric fields for matched stimulation conditions across species.

There are several key findings regarding TMS: (i) the electric field strength first increases with increasing head size and then decreases; (ii) the deflection point of this function depends on the coil size with the smaller coil showing a decrease in electric field strength starting at a smaller head size; (iii) the relative affected brain volume decreases with head size; (iv) the tangential electric field component dominates the perpendicular component across all species, but more so in the mouse model. For the 70 mm figure-8 coil, the maximum induced electric field is 55% lower in the mouse than in the monkey and human, where the field strengths are comparable. The use of a smaller 25 mm figure-8 coil leads to comparable electric fields in the mouse and human, but 30% stronger electric field in the monkey. Both coils affect a smaller relative brain volume in the human in comparison to the monkey (~ 4-times) and mouse (~ 20-times).

Analyzing TMS-induced electric fields in anatomically realistic and simplified spherical models, we found an increase in electric field strength with head size which was reversed to a decrease for even larger radii. A possible mechanism for this effect is that the head underneath the TMS coil captures a fraction of the total magnetic flux. This fraction increases with increasing head size; thus, the induced electric field gets stronger (Weissman et al., 1992). However, once the head reaches a certain dimension relative to the coil size, the captured energy is maximized, but the induced electric current is spread through a larger conductive volume and thus creates a weaker electric field. For a smaller TMS coil, the maximum magnetic flux captured occurs at a smaller head size, which results in an earlier decrease.

Our key findings for TES are: (i) the electric field strength in the brain decreases with increasing head size; (ii) the electric field strength and affected brain volume strongly vary with the electrode montage; (iii) the tangential field is larger than the perpendicular component in all models, but more so in the mouse. We attribute the strong decrease of the TES electric field with increasing head size, which is notably opposite to the relationship found in TMS, to progressively thicker layers of tissues around the brain and thus increasingly diluted current density. The poorly conducting skull isolates the brain, while highly conductive soft tissues and CSF provide avenues for current shunting. This resulted in up to 100-times higher electric fields in the mouse model compared to the human model for the same stimulation intensity. Electric fields in the monkey model were found to be ~ 3-times higher than in the human. On average, the stimulation affects a 1.9-times larger brain volume in the monkey and a 2.5-times larger relative brain volume in the mouse than in the human. Of note is the high variability of electric field strengths for different electrode montages. Thus, for every given experiment, it will be important to consider both the electrode montage and individual anatomy.

Both TMS and TES simulations indicate that specific properties of the induced electric fields in the human brain are better captured in monkeys than in mice. Besides a more comparable electric field strength in the brain and affected brain volume, the ratio of tangential to perpendicular electric field components is similar in human and monkey models. In the mouse, the tangential field dominates (~ 60-80% higher) for every modality and condition. While the electric field strength can be easily scaled (within safety limits) by adjusting the TMS or TES intensity, the ratio of electric field components cannot. The difference across species is due to the pronounced gyrification of the primate brains in comparison to rodents. However, the implications of these different electric field components on the resulting physiological effects are less clear. It is known that the electric field predominantly affects neural cells that are oriented parallel to the electric field (Aspart et al., 2018; Radman et al., 2009; Rawji et al., 2018; Terzuolo and Bullock, 1956), and a critical mass of affected neurons is necessary to elicit a system-level response. The dissimilar balance of tangential to perpendicular electric field components in the mouse brain could change the system-level response to NIBS relative to the human. Although this is speculative, research in non-human primates should remain essential for understanding the mechanisms of brain stimulation.

In this study, we provide a comprehensive evaluation of TMS and TES electric fields across different species. Our results validate and significantly expand on previous modeling efforts. One earlier study investigated the TMS-induced electric field with respect to brain size using spherical models (Weissman et al., 1992). It demonstrated a steady up to 5-times increase in the induced electric field magnitude as a function of increasing model radius from 0.5 to 7.5 cm, which agrees with our results for the given size range and a large coil. We also confirm a broad TMS-induced electric field in the mouse brain, which is qualitatively different from the one in humans (Crowther et al., 2014; Salvador and Miranda, 2009). While using a smaller TMS coil reduces the relative stimulation volume in rodents, it is still much larger than in humans. Intracranial application of short-pulsed electric currents might be a way to mimic the TMS-induced electric field in a mouse in a realistic manner (Barnes et al., 2014). Electric fields in humans and monkeys due to TMS using a standard 70 mm figure-8 coil are largely similar. An important caveat is that this was shown here with one specific human and monkey model and can slightly differ for other individuals in a given experiment.

A previous modelling study of TES in a mouse with a bihemispheric electrode montage showed a maximum electric field strength of ~ 20 mV/mm per 1 mA in the brain (Bernabei et al., 2014), which is in good agreement with our results. This led researchers to assume a ratio of 50:1 in the TES electric field in mouse relative to humans, where we would expect a maximum field strength of ~ 0.4 mV/mm (Huang et al., 2017; Opitz et al., 2016). However, we demonstrate that other electrode montages can create stronger electric fields, up to 50 mV/mm. On average, our estimate of electric field strengths in a mouse model roughly relate to those in human as 100:1. Thus, a typical TES intensity of 1-2 mA in humans (Antal et al., 2017; Paulus et al., 2016) approximately translates to mice as 10-20 μA and monkeys as 0.3-0.6 mA. Commonly used intensities of 0.1 mA in previous studies in mice (e.g., Grossman et al., 2017; Monai et al., 2016) lead to electric field strengths that are way above what is safe and tolerable in human application (Nitsche and Bikson, 2017). Importantly, TES electric fields in animals and humans strongly depend on the specific electrode montage and individual anatomy. Thus, precise measurements and calculations for every specific case can improve the transferability of results. Nevertheless, our modeling results of intracranial electric fields are in good agreement with existing *in vivo* measurements in monkeys and humans (Alekseichuk et al., 2018; Huang et al., 2017; Opitz et al., 2018, 2016).

In conclusion, we provide a systematic evaluation of TES- and TMS-induced electric fields in two popular animal models and compare them to the human case. We outline differences and similarities between electric fields across species and draw attention to the effects of head/brain size and brain gyrification. Notably TMS and TES differ in their relationship of electric field strength and head size in almost an opposite manner. To generate the same intracranial electric field strength in a mouse as in a human, TMS intensity should typically be *higher*, yet TES intensity should be two orders of magnitude *lower.* For a monkey, our data advocate the use of the *same* TMS intensity and three times *lower* TES intensity to what is applicable in humans. However, while we provide general guidelines for scaling TMS and TES stimulation parameters across species, significant variability in the electric field strength across individuals and stimulation montages stresses the importance of exact estimations for every experiment and individual participant.

## Acknowledgments

This work was supported by the University of Minnesota’s MnDRIVE Initiative, the Institute for Engineering in Medicine at the University of Minnesota, and NIH grant R01MH118930.

## Appendix A. Data tables

**Table A1.** Summary of TMS modeling results (70 mm figure-8 coil). The following coil orientations targeting the left central brain area (“motor cortex”) were simulated for the mouse, monkey, and human models: caudal-medial (CM), medial (M), and caudal-lateral (CL). The affected area L_½max_ is defined as the volume or surface where the electric field strength is equal or greater than the half-maximum for a given simulation. The robust maximum E_max_ corresponds to the 99.9 percentile of the simulated electric field strength in the volume or on the surface.

| TMS | Mouse |  |  | Capuchin Monkey |  |  | Human |  |  |
| --- | --- | --- | --- | --- | --- | --- | --- | --- | --- |
|  | CM | M | CL | CM | M | CL | CM | M | CL |
| Grey Matter Volume |  |  |  |  |  |  |  |  |  |
| $E_{max}$, mV/mm | 59.13 | 51.62 | 59.01 | 128.35 | 125.59 | 125.38 | 124.58 | 123.37 | 132.33 |
| $E_{median}$, mV/mm | 26.16 | 20.49 | 23.57 | 28.83 | 24.13 | 25.84 | 11.61 | 10.34 | 11.43 |
| $L_{\frac{1}{2}max}$, % | 40.97 | 32.67 | 33.72 | 8.65 | 8.12 | 9.50 | 2.00 | 2.66 | 2.19 |
| Grey Matter Surface |  |  |  |  |  |  |  |  |  |
| $E_{max}$, mV/mm | 62.46 | 53.18 | 60.42 | 144.21 | 138.73 | 134.39 | 134.22 | 133.09 | 139.82 |
| $E_{median}$, mV/mm | 26.31 | 21.08 | 23.84 | 27.18 | 22.00 | 24.14 | 12.70 | 11.75 | 12.62 |
| $L_{\frac{1}{2}max}$, % | 40.60 | 37.03 | 37.93 | 6.63 | 6.59 | 8.37 | 1.93 | 2.58 | 2.21 |

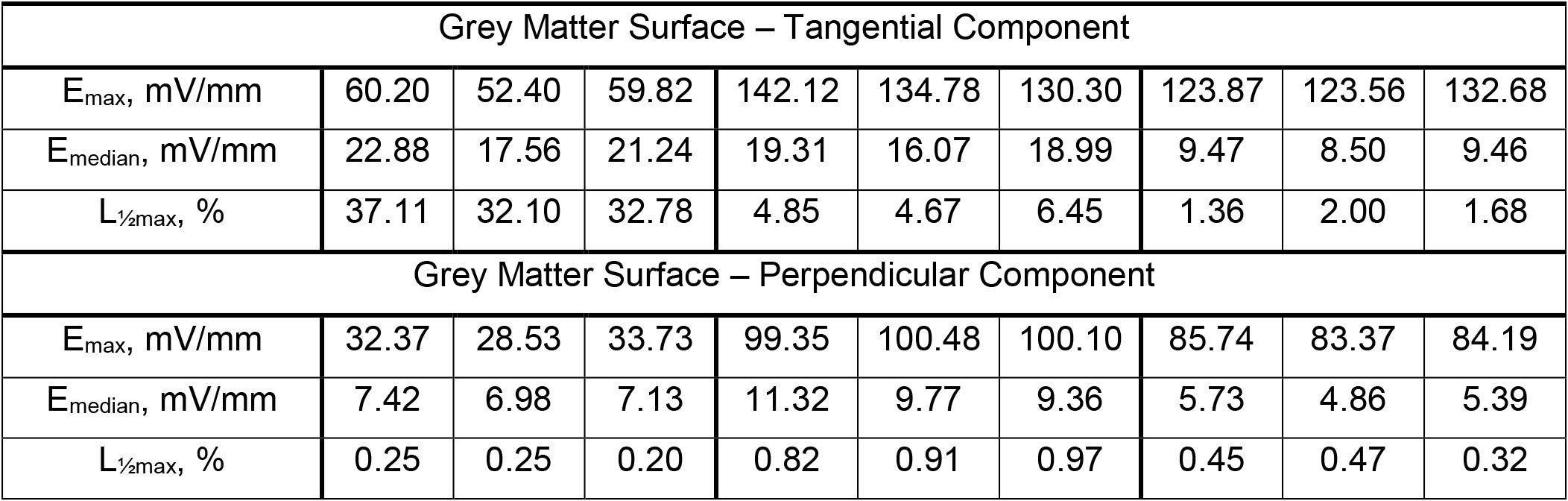

| Grey Matter Surface – Tangential Component |  |  |  |  |  |  |  |  |  |
| --- | --- | --- | --- | --- | --- | --- | --- | --- | --- |
| $E_{\max}$, mV/mm | 60.20 | 52.40 | 59.82 | 142.12 | 134.78 | 130.30 | 123.87 | 123.56 | 132.68 |
| $E_{\text{median}}$, mV/mm | 22.88 | 17.56 | 21.24 | 19.31 | 16.07 | 18.99 | 9.47 | 8.50 | 9.46 |
| $L_{\frac{1}{2}\max}$, % | 37.11 | 32.10 | 32.78 | 4.85 | 4.67 | 6.45 | 1.36 | 2.00 | 1.68 |
| Grey Matter Surface – Perpendicular Component |  |  |  |  |  |  |  |  |  |
| $E_{\max}$, mV/mm | 32.37 | 28.53 | 33.73 | 99.35 | 100.48 | 100.10 | 85.74 | 83.37 | 84.19 |
| $E_{\text{median}}$, mV/mm | 7.42 | 6.98 | 7.13 | 11.32 | 9.77 | 9.36 | 5.73 | 4.86 | 5.39 |
| $L_{\frac{1}{2}\max}$, % | 0.25 | 0.25 | 0.20 | 0.82 | 0.91 | 0.97 | 0.45 | 0.47 | 0.32 |

**Table A2.** Summary of TMS modeling results (25 mm figure-8 coil). The following coil orientations targeting the left central brain area (“motor cortex”) were simulated for the mouse, monkey, and human models: caudal-medial (CM), medial (M), and caudal-lateral (CL). The affected area L_½max_ is defined as the volume or surface where the electric field strength is equal or greater than the half-maximum for a given simulation. The robust maximum E_max_ corresponds to the 99.9 percentile of the simulated electric field strength in the volume or on the surface.

| TMS | Mouse |  |  | Capuchin Monkey |  |  | Human |  |  |
| --- | --- | --- | --- | --- | --- | --- | --- | --- | --- |
|  | CM | M | CL | CM | M | CL | CM | M | CL |
| Grey Matter Volume |  |  |  |  |  |  |  |  |  |
| $E_{\max}$, mV/mm | 84.24 | 75.38 | 84.25 | 118.43 | 113.12 | 108.53 | 83.88 | 84.22 | 91.31 |
| $E_{\text{median}}$, mV/mm | 35.66 | 29.10 | 32.24 | 20.57 | 17.32 | 17.85 | 4.34 | 3.78 | 4.20 |
| $L_{\frac{1}{2}\max}$, % | 37.99 | 30.82 | 31.57 | 5.84 | 5.64 | 6.66 | 1.28 | 1.52 | 1.09 |
| Grey Matter Surface |  |  |  |  |  |  |  |  |  |
| $E_{\max}$, mV/mm | 89.39 | 77.81 | 85.74 | 134.53 | 126.72 | 119.25 | 94.25 | 96.28 | 99.93 |
| $E_{\text{median}}$, mV/mm | 35.68 | 29.77 | 32.23 | 19.57 | 16.18 | 16.59 | 4.78 | 4.33 | 4.63 |
| $L_{\frac{1}{2}\max}$, % | 37.66 | 34.58 | 36.14 | 4.46 | 4.69 | 5.91 | 1.16 | 1.34 | 1.10 |
| Grey Matter Surface – Tangential Component |  |  |  |  |  |  |  |  |  |
| $E_{\max}$, mV/mm | 85.96 | 76.66 | 85.35 | 131.80 | 122.42 | 114.84 | 85.64 | 87.54 | 94.54 |
| $E_{\text{median}}$, mV/mm | 30.80 | 24.20 | 28.65 | 13.63 | 11.86 | 12.59 | 3.58 | 3.13 | 3.51 |
| $L_{\frac{1}{2}\max}$, % | 35.16 | 30.33 | 31.01 | 3.36 | 3.53 | 4.94 | 0.88 | 1.08 | 0.80 |
| Grey Matter Surface – Perpendicular Component |  |  |  |  |  |  |  |  |  |
| $E_{\max}$, mV/mm | 45.32 | 40.60 | 46.74 | 90.30 | 84.37 | 81.44 | 56.64 | 55.75 | 56.11 |
| $E_{\text{median}}$, mV/mm | 10.22 | 10.23 | 9.94 | 8.33 | 6.74 | 6.39 | 2.17 | 1.79 | 2.03 |
| $L_{\frac{1}{2}\max}$, % | 0.19 | 0.19 | 0.16 | 0.63 | 0.61 | 0.57 | 0.28 | 0.24 | 0.21 |

**Table A3.**
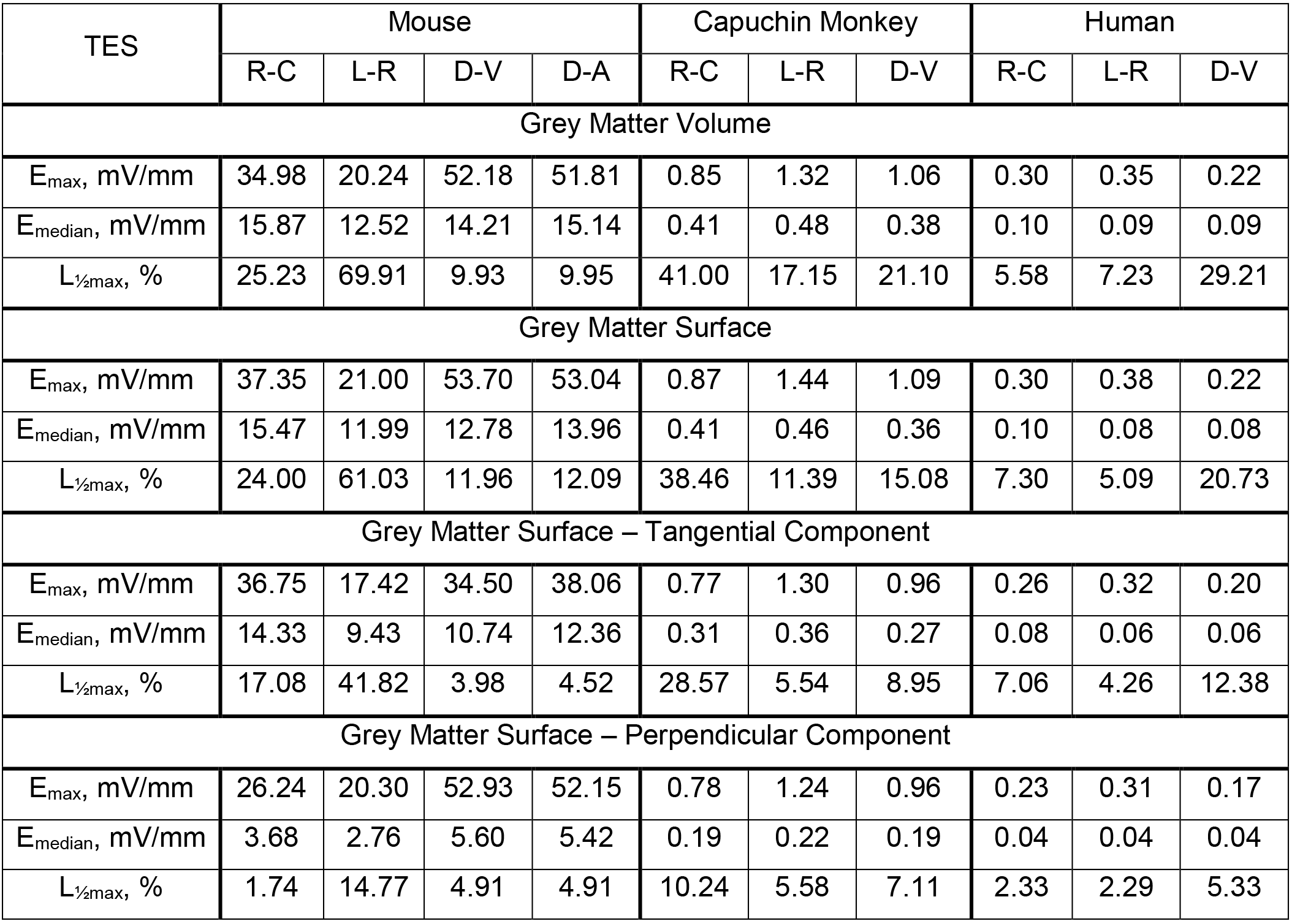
Summary of TES modeling results. The following montages were simulated for the mouse, capuchin monkey, and human model: rostral-caudal (R-C), left-right (L-R), and dorsal-ventral (D-V). In addition, a dorsal-abdominal (D-A) montage is included for the mouse model. The affected area L_½max_ is defined as the volume or surface where the electric field strength is equal or greater than the half-maximum for a given simulation. The robust maximum E_max_ corresponds to the 99.9 percentile of the simulated electric field strength in the volume or on the surface.

**Table A4.** Summary of TES modeling. The following two-electrode montages with variable electrode sizes are estimated for the human model: rostral-caudal (R-C), left-right (L-R), and dorsal-ventral (D-V). The affected area L_½max_ is defined as the volume or surface where the electric field strength is equal or greater than the half-maximum for a given simulation. The robust maximum E_max_ corresponds to the 99.9 percentile of the simulated electric field strength in the volume or on the surface.

| TES | Electrode d = 36 mm |  |  | Electrode d = 15 mm |  |  | Electrode d = 3 mm |  |  |
| --- | --- | --- | --- | --- | --- | --- | --- | --- | --- |
|  | R-C | L-R | D-V | R-C | L-R | D-V | R-C | L-R | D-V |
| Grey Matter Volume |  |  |  |  |  |  |  |  |  |
| $E_{\max}$, mV/mm | 0.30 | 0.35 | 0.22 | 0.32 | 0.40 | 0.21 | 0.31 | 0.43 | 0.22 |
| $E_{\text{median}}$, mV/mm | 0.10 | 0.09 | 0.09 | 0.10 | 0.09 | 0.09 | 0.09 | 0.09 | 0.09 |
| $L_{\frac{1}{2}\max}$, % | 5.58 | 7.23 | 29.21 | 3.53 | 5.28 | 29.57 | 3.30 | 4.77 | 29.70 |

| Grey Matter Surface |  |  |  |  |  |  |  |  |  |
| --- | --- | --- | --- | --- | --- | --- | --- | --- | --- |
| $E_{\max}$, mV/mm | 0.30 | 0.38 | 0.22 | 0.32 | 0.44 | 0.21 | 0.31 | 0.46 | 0.22 |
| $E_{\text{median}}$, mV/mm | 0.10 | 0.08 | 0.08 | 0.10 | 0.08 | 0.08 | 0.09 | 0.09 | 0.08 |
| $L_{\frac{1}{2}\max}$, % | 7.30 | 5.09 | 20.73 | 5.03 | 3.62 | 21.16 | 4.78 | 3.34 | 21.25 |
| Grey Matter Surface – Tangential Component |  |  |  |  |  |  |  |  |  |
| $E_{\max}$, mV/mm | 0.26 | 0.32 | 0.20 | 0.27 | 0.37 | 0.20 | 0.26 | 0.39 | 0.20 |
| $E_{\text{median}}$, mV/mm | 0.08 | 0.06 | 0.06 | 0.08 | 0.07 | 0.06 | 0.08 | 0.07 | 0.06 |
| $L_{\frac{1}{2}\max}$, % | 7.06 | 4.26 | 12.38 | 5.34 | 3.00 | 12.73 | 5.09 | 2.78 | 12.85 |
| Grey Matter Surface – Perpendicular Component |  |  |  |  |  |  |  |  |  |
| $E_{\max}$, mV/mm | 0.23 | 0.31 | 0.17 | 0.24 | 0.35 | 0.17 | 0.24 | 0.37 | 0.17 |
| $E_{\text{median}}$, mV/mm | 0.04 | 0.04 | 0.04 | 0.04 | 0.04 | 0.04 | 0.04 | 0.04 | 0.04 |
| $L_{\frac{1}{2}\max}$, % | 2.33 | 2.29 | 5.33 | 1.93 | 1.63 | 5.44 | 1.87 | 1.52 | 5.47 |

## Appendix B. Analytical formulation of TMS-induced electric field

The analytical solution of the TMS-induced electric field follows the method by Eaton derived for a homogeneous spherical volume conductor for an arbitrary TMS coil geometry (Eaton, 1992). Below we summarize the essential equations and their derivation for a single loop figure-8 coil. To simplify the equations, the origin of the spherical coordinate system is set as the center of the model. Time varying signals have the form *e*^*jωt*^, where 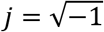 and ω is the angular frequency. The electric field ***E*** first depends on the complex vector constant *C*_*lm*_ that relates to a given coil geometry and placement:

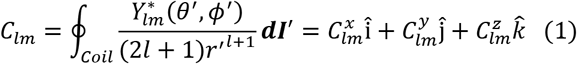

Where * is the complex conjugate; *r*′, *θ*′, *ϕ*′ are radial distance, polar angle, and azimuthal angle of the differential segment of the coil, *Y*_*lm*_(*θ*′, *ϕ*′) are spherical harmonic functions; *I* is the current in the coil; ***dl***′ is the orientation of the current in the coil along the current path. Further we define *D*_*lm*_, *E*_*lm*_, and *F*_*lm*_ to simplify the equations:

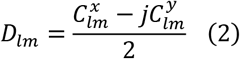

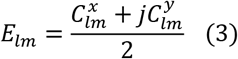

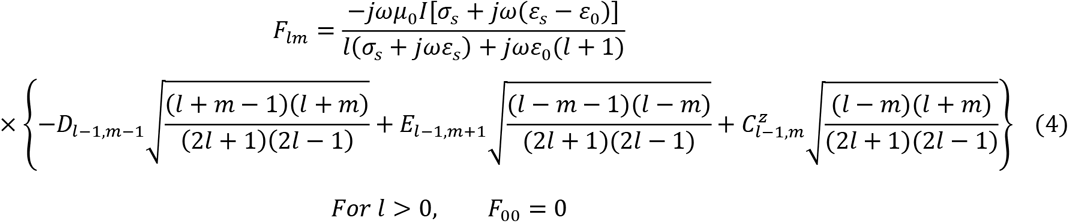

We assume that the sphere has the permeability of free space *μ*_0_, where ε_*s*_ and σ_*s*_ are the permittivity and conductivity of the homogenous isotropic sphere, respectively. We used ε_*s*_ ≈ 13000ε_*0*_ and σ_*s*_ = 0.14 *sm*^−1^. Using the equations above, the electric field at any given point in space (*r*,*θ*,*ϕ*) along three spherical axes is:

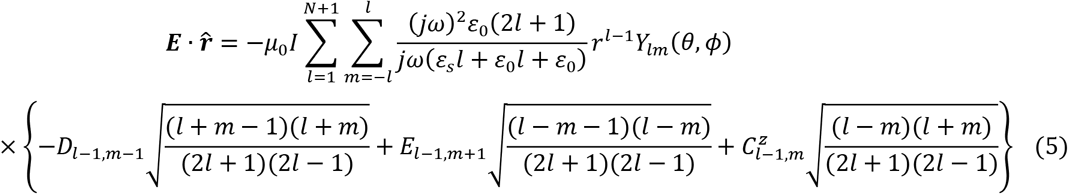

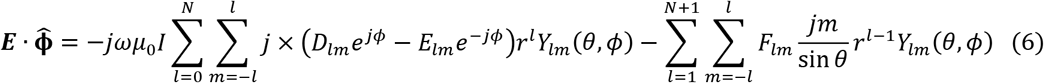

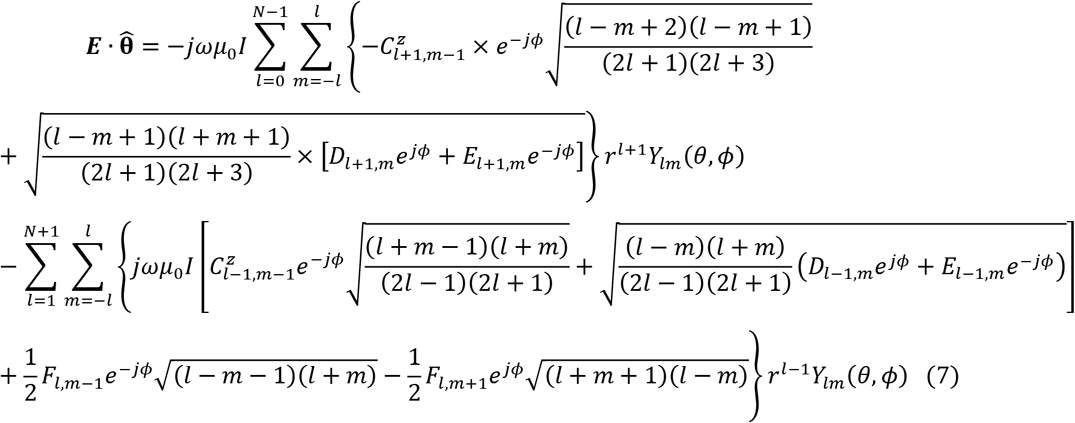

Equations 5-7 are N^th^ order approximations of the analytical solution. The result converges to the exact solution with *N* → ∞. In our calculations, we used N = 20 which gives sufficient accuracy based on the convergence rate. In what follows, we define *C*_*lm*_ for a one wire loop figure-8 coil (Figure B1).

**Figure B1.**
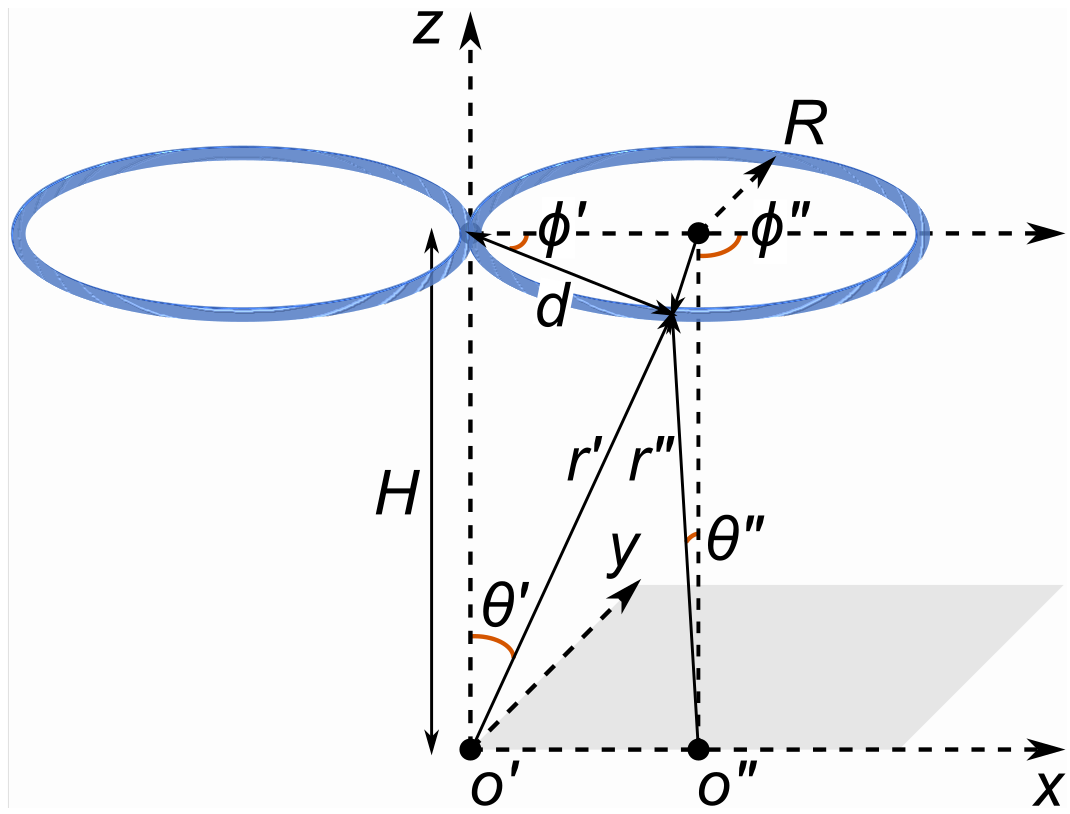
One wire loopfigure-8 coil in a spherical coordinate system.

The coil with radius *R* is parallel to the x-y plane, with the loops along the x-axis, and the center of the coil being a distance *H* (in our case, 4 mm) above the head. *C*_*lm*_ is calculated for both loops of the coil separately. For the loop on the positive side of the x-axis, to simplify the calculations, we translated the coordinate system in such way that the center of the loop is directly above the new center of the coordinate system. The old coordinate system relates to the new coordinate system as follows:

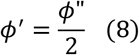

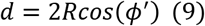

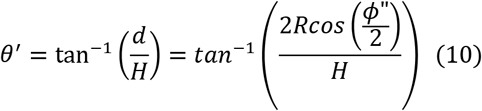

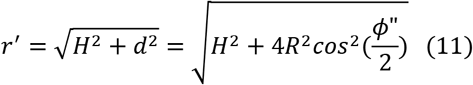

Given such coordinate translation and a counterclockwise direction of coil currents, the integral can be written as:

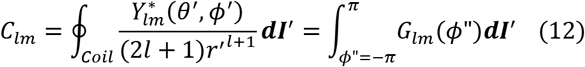

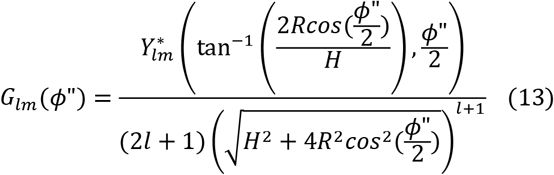

Therefore, by expanding the integral over x, y and z orientations, we arrive at:

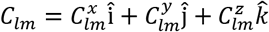

Additionally, we know that in the new coordinate system

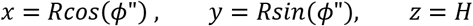

Then,

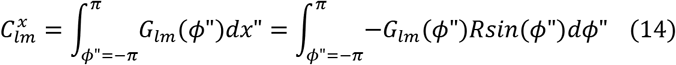

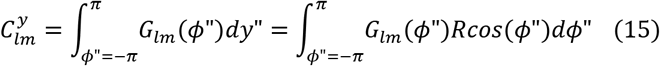

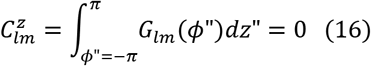

All the above steps can be repeated for the other side of the loop with the opposite current direction. The only change will be:

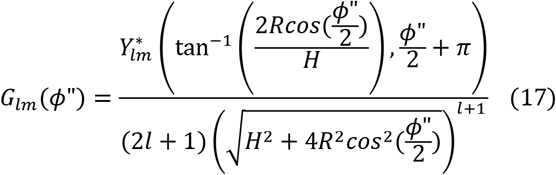

Finally, both integral results are added together to calculate *C*_*lm*_ for the whole figure-8 coil.

## Appendix C. Supplementary data

Supplementary materials associated with this article can be found below.

## SUPPLEMENTARY MATERIALS

**Supplementary Figure 1.**
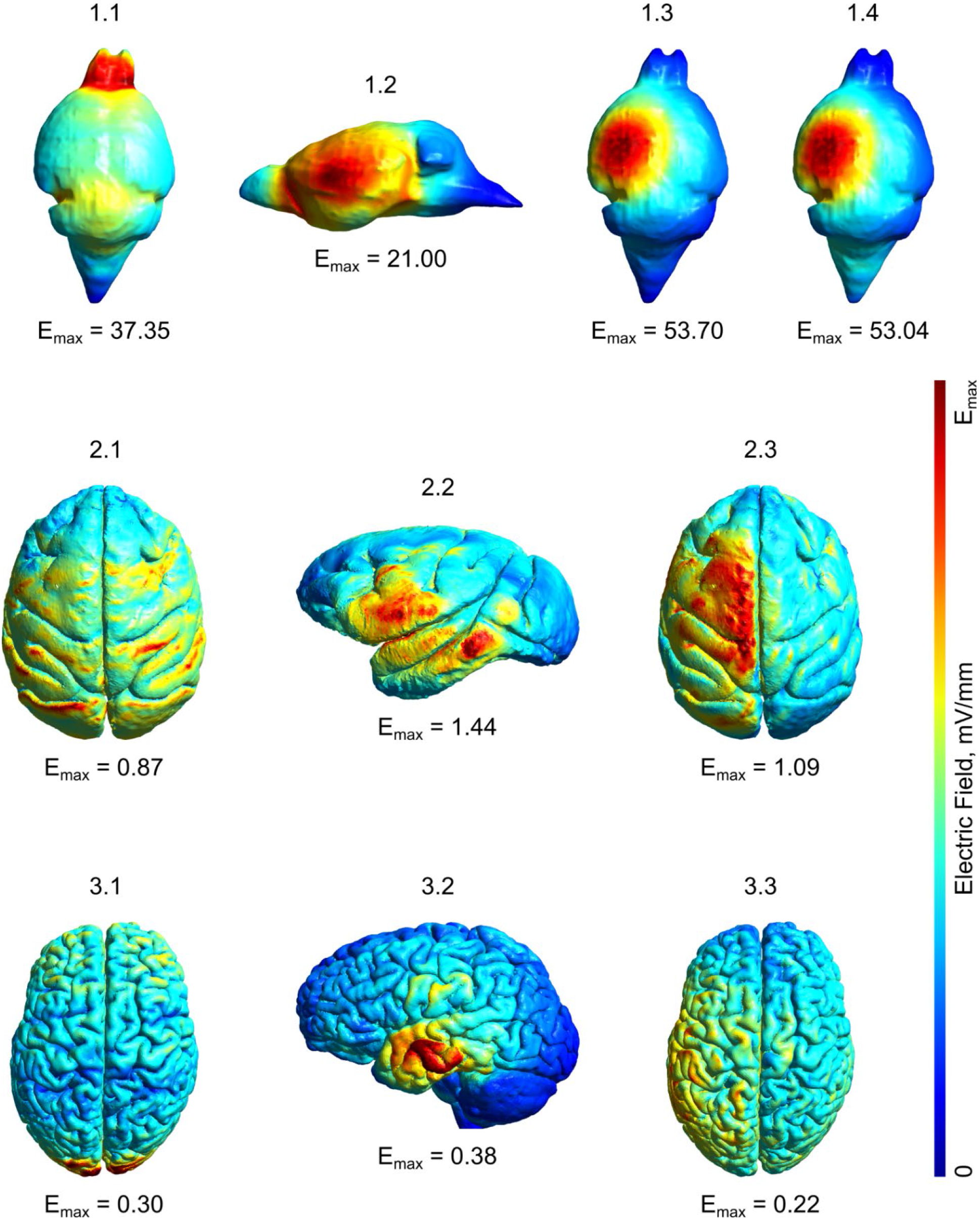
TMS electric fields for the 70 mm figure-8 coil in the (1.x) mouse, (2.x) monkey, and (3.x) human models. The coil is located above the left central brain area (“motor cortex”) and rotated towards (x.1) caudal medial, (x.2) medial, or (x.3) caudal lateral ends.

**Supplementary Figure 2.**
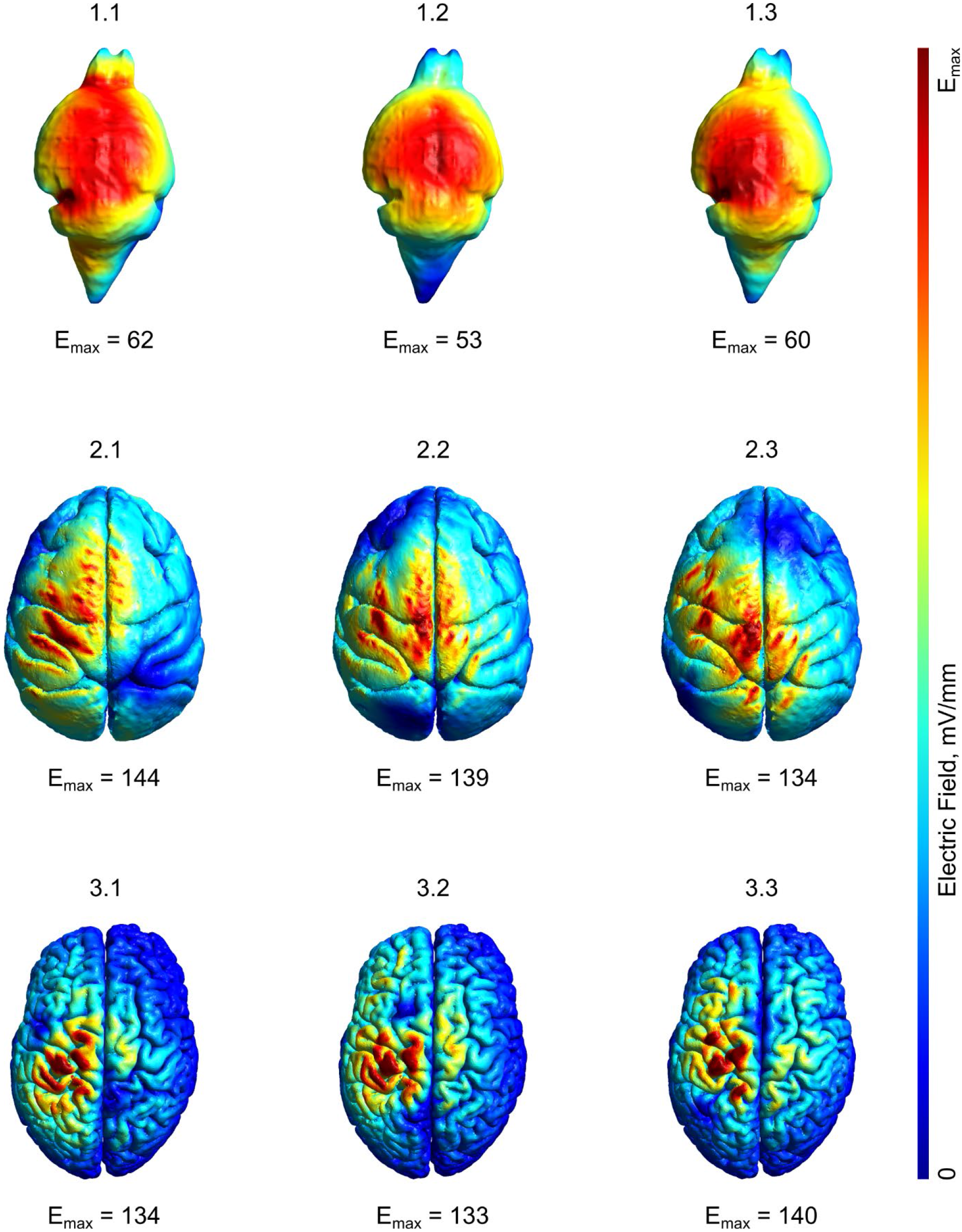
TMS electric fields for the 25 mm figure-8 coil in the (1.x) mouse, (2.x) monkey, and (3.x) human models. The coil is located above the left central brain area (“motor cortex”) and rotated towards (x.1) caudal medial, (x.2) medial, or (x.3) caudal lateral ends.

**Supplementary Figure 3.**
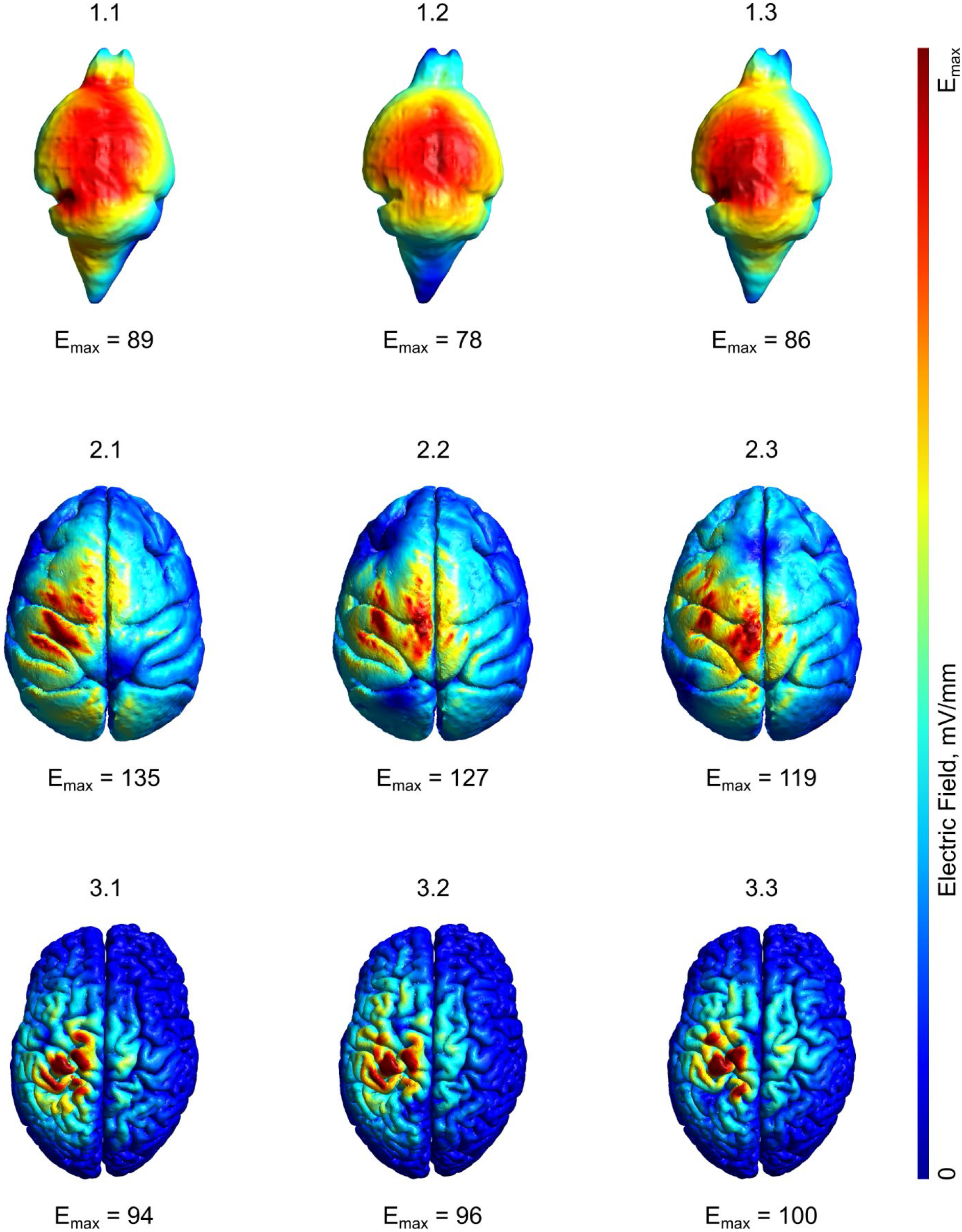
TES electric fields for different montages in the (1.x) mouse, (2.x) monkey, and (3.x) human models. The electrode locations are the following: x.1 – medial rostral and caudal brain areas (R-C), x.2 – central left and right brain areas (L-R), x.3 – left central brain area and right shoulder/neck area (D-V), and x.4 – left dorsal brain area and abdominal area (D-A).

**Supplementary Figure 4.**
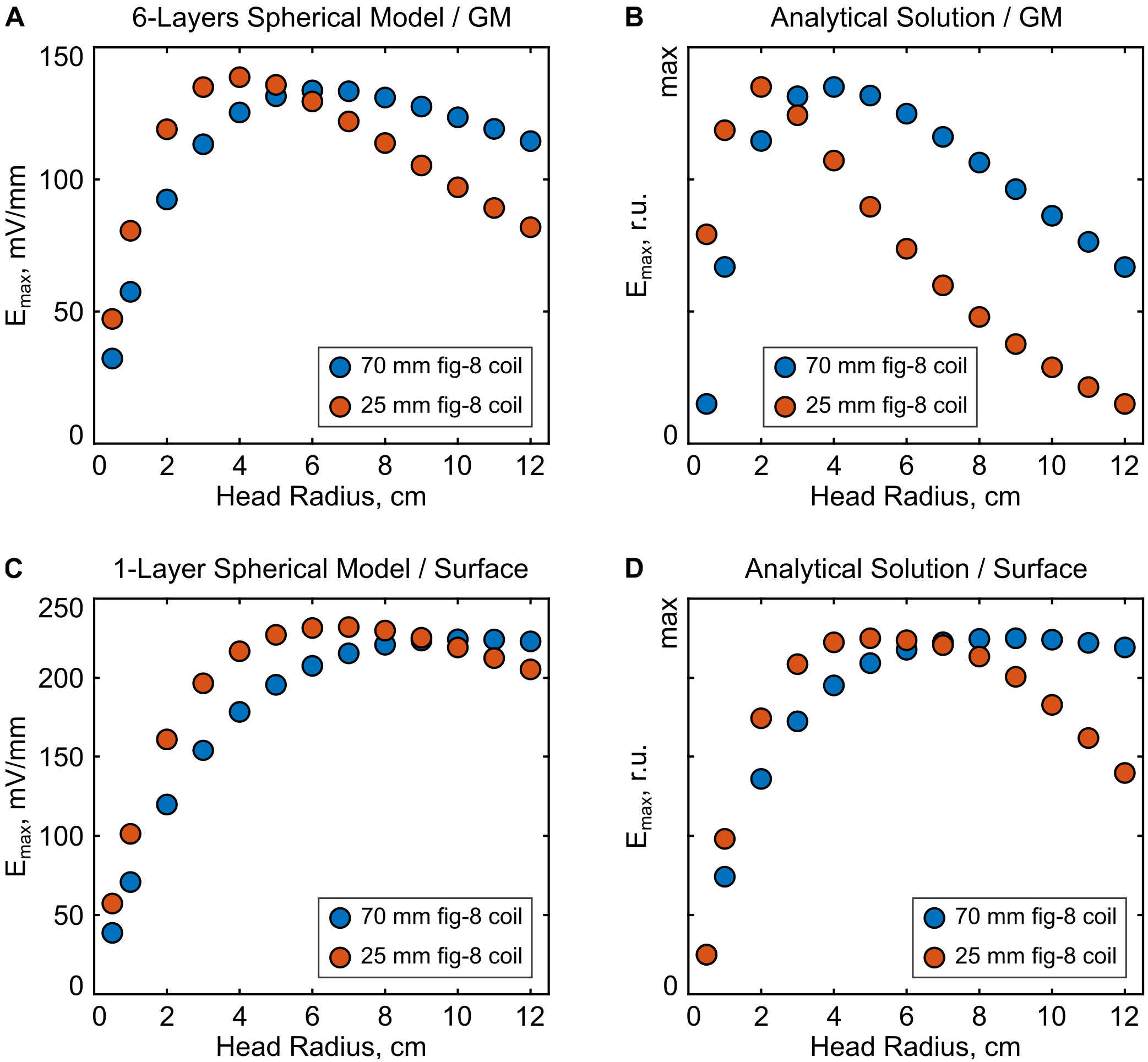
TMS-induced electric field strength as a function of head size using spherical models. Results are shown both using the FEM simulation approach (A, C) and an analytical solution (B, D). Note that the TMS coils and head models were not identical between the numerical and analytical solution which results in slightly varying results. Top row: Electric field strength in the “grey matter”, which accounts for changes in the coil-to-brain distance with increasing head size. Bottom row: results on the outer spherical surface, which represent the same coil-to-brain distance for every model size.

**Supplementary Figure 5.**
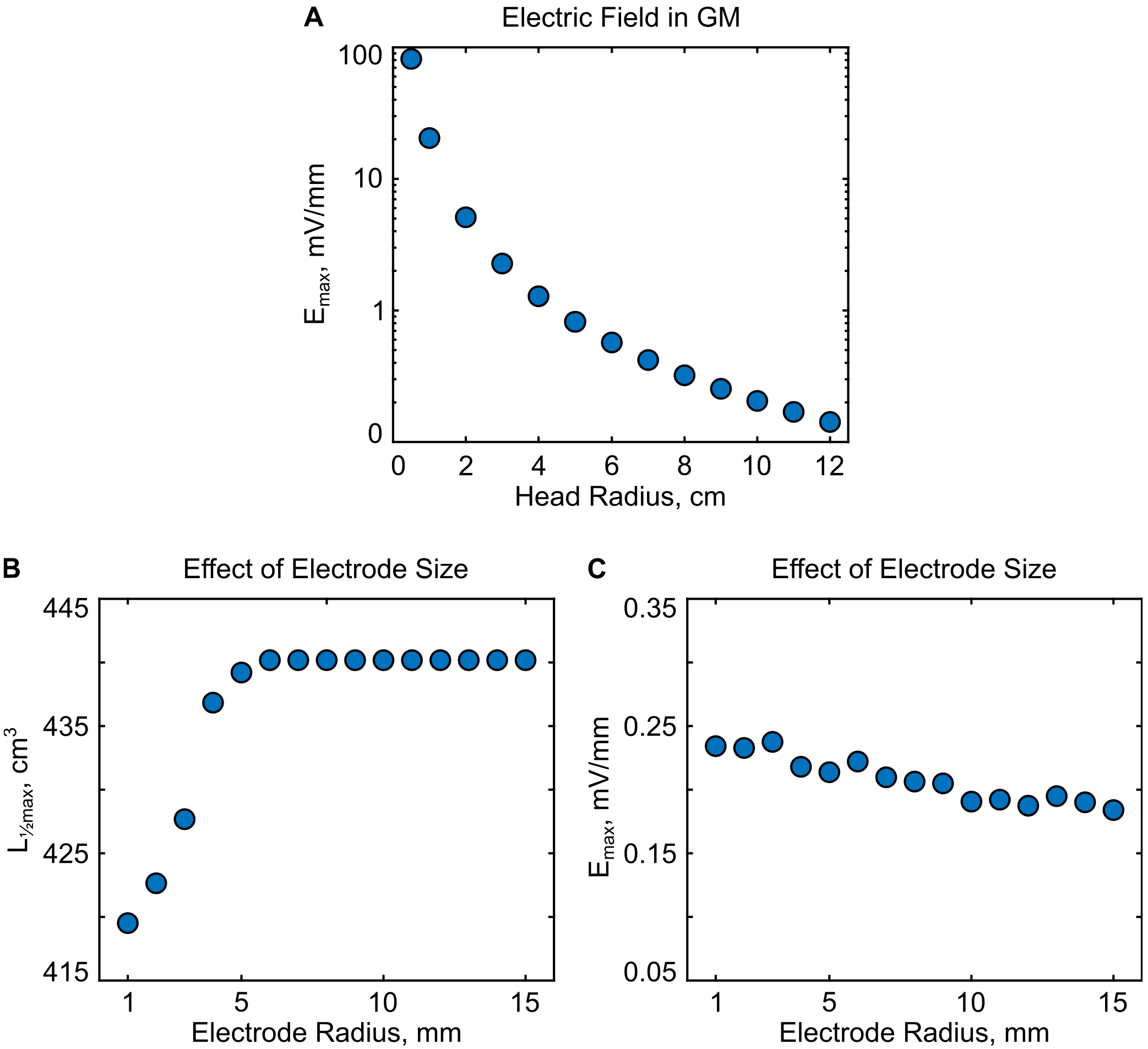
(A) TES electric field in the “grey matter” for a six-layer spherical model. (B-C) The effect of the stimulation electrode size on the affected volume (B) and electric field strength (C) in the “grey matter” of the spherical model with r = 10 cm, which approximately corresponds to human head size.

